# SPRDA: a matrix completion approach based on the structural perturbation to infer disease-associated Piwi-Interacting RNAs

**DOI:** 10.1101/2020.07.02.185611

**Authors:** Kai Zheng, Zhu-Hong You, Lei Wang, Leon Wong, Zhao-hui Zhan

## Abstract

Emerging evidence suggests that PIWI-interacting RNAs (piRNAs) are one of the most influential small non-coding RNAs (ncRNAs) that regulate RNA silencing. piRNA and PIWI proteins have been confirmed for disease diagnosis and treatment as novel biomarkers due to its abnormal expression in various cancers. However, the current research is not strong enough to further clarify the functions of piRNA in cancer and its underlying mechanism. Therefore, how to provide large-scale and serious piRNA candidates for biological research has grown up to be a pressing issue. The main motivation of this work is tantamount to fill the gap in research on large-scale prediction of disease-related piRNAs. In this study, a novel computational model based on the structural perturbation method is proposed, called SPRDA. In detail, the duplex network is constructed based on the piRNA similarity network and disease similarity network extracted from piRNA sequence information, Gaussian interaction profile kernel similarity information and gene-disease association information. The structural perturbation method is then used to predict the potential associations on the duplex network, which is more predictive than other network structures in terms of structural consistency. In the five-fold cross-validation, SPRDA shows high performance on the benchmark dataset piRDisease, with an AUC of 0.9529. Furthermore, the predictive performance of SPRDA for 10 diseases shows the robustness of the proposed method. Overall, the proposed approach can provide unique insights into the pathogenesis of the disease and will advance the field of oncology diagnosis and treatment.

## 1 Introduction

PIWI interacting RNA (piRNA) is a distinct class of small non-coding RNA mainly expressed in germ cells [1, 2]. In 2006, Girard *et al*. discovered a previously uncharacteristic class of small RNAs that bind to murine Piwi proteins MIWI and are highly abundant in the testes. Therefore, these small RNAs are named Piwi-interacting RNAs (piRNAs) [3]. piRNA specifically binds to the PIWI subfamily of Argonaute proteins to form an effector complex to silence transposon activity in animal gonads, and maintain germline genome integrity through transcription or post-transcription mechanisms [4-7]. In addition, studies have demonstrated that piRNA can also regulate cellular genes and transmit “self” and “non-self” memories during intergenerational inheritance, which implicates PIWI in establishment of transgenerational epigenetic states [8-10]. In metazoans, there are 3 types of small silencing RNA: microRNA (miRNA), piRNA and small interfering RNA (siRNA). The processing of both miRNAs and siRNAs need to cleave by Dicer endonuclease. Compare with other small RNAs, piRNAs arise by Dicer-independent mechanisms [9, 11-13]. Different biogenesis mechanisms led to the realization that piRNA is manifest from siRNA and miRNA.

As research continues, piRNA and PIWI proteins have been confirmed for disease diagnosis and treatment as novel biomarkers due to its abnormal expression in various cancers [14-19]. Therefore, exploring the mechanism of piRNA at the transcriptional or post-transcriptional level in pathological processes has become an important way to figure out the molecular roots of human disease. For example, Li *et al*. found that piR-32051 was positively correlated with high tumor stage and metastasis of Kidney cancer [20]. Subsequently, Mark *et al*. confirmed that the expression of piR-34871 and piR-52200 was positively correlated with RASSF1C expression in tumor tissue. In addition, knock-down of piR-34871 and piR-52200 was in a position to significantly reduce the proliferation of lung cancer cell lines (A549 and H1299) in 2017 [21]. In addition, Anna *et al*. Found that piR-651 expression was inhibited in serum samples from 94 Hematological malignancy patients. Interestingly, after complete remission, piR-651 levels returned to normal [22]. The piR-39980 in fibrosarcoma was first reported in 2019. Das *et al*. found that transient overexpression of piR-39980 can attenuate the proliferation and induce apoptosis of HT1080 fibrosarcoma cells, and has a strong antitumor effect [23]. Glioblastoma has also been linked to piRNA, Leng *et al*. found that overexpression of piR-DQ590027 increased the permeability of the blood-brain barrier under glioma conditions and promoted the transport of anti-glioma drugs to glioma tissue [24].

piRNA is accumulating rapidly as research continues. In order to homogenize complex and heterogeneous data, some databases were established to collect the features of piRNA such as piRNAQuest [25], piRNABank [26] and piRBase [27]. In 2019, Muhammad *et al*. developed the piRDisease database in order to promote research on piRNAs in disease diagnosis, prognosis, and assessment of treatment response [28]. The database provides comprehensive and proprietary data, including experimental support, sequence and location information. Although many predictors have been proposed to infer the link between ncRNA and disease, piRNA-based calculation model is unexplored [29-33].

In this study, a piRNA-disease association predictor based on the structural perturbation method was proposed, called SPRDA. The SPRDA has following advances: (i) We construct a duplex network based on the piRNA similarity network and disease similarity network extracted from piRNA sequence information and gene-disease association information, which can effectively improve prediction performance. (ii) A new uniquely mapped disease characterization method is proposed and used to quantify the similarity. The results demonstrate that the method has lower computational cost under the same performance. (iii) We use of the structural consistency index to quantify the link predictability of the piRNA-disease network. The results show that the network consistency is much higher than other comparative networks. In the 5-fold cross-validation, the AUC of our method reaches 91.45% and the accuracy is 84.49%. In addition, we performed independent performance tests on 12 different diseases to evaluate the performance of SPRDA. From the results, it can effectively identify disease-related piRNAs. Overall, the proposed method can provide unique insights into the mechanisms of disease and will further the field of tumor diagnosis and treatment.

## 2 Materials and Methods

### 2.1 Benchmark dataset

Many public databases provide piRNA feature information, such as piRNAQuest [25], piRNABank [26] and piRBase [27]. However, only piRDisease [28] currently contains experimentally verified piRNA-disease associations. To enable large-scale analysis of disease-related piRNAs, we used piRDisease as the baseline dataset (http://www.piwirna2disease.org/index.php). Since there are a large number of nodes with *degree* = 1 in the piRNA-disease association network formed by this database, which seriously affects the predictability of the network.Therefore, we performed preprocessing on the original data by deduplication and filtering out points with *degree* = 1, and define the benchmark data set piRD, as shown in Table 1.

**Table 1.** The details of benchmark dataset piRD.

| Benchmark dataset | piRNAs | diseases | Associations |
| --- | --- | --- | --- |
| piRD | 501 | 22 | 1212 |

### 2.2 Construct piRNA–disease bilayer network

The piRNA–disease duplex network is constructed by three networks, including the disease similarity network, piRNA similarity network, and piRNA-disease association network. The disease similarity network and piRNA similarity network are weighted by different subnets.

#### The piRNA similarity network

The piRNA sequence similarity network and the piRNA Gaussian interaction profile kernel similarity network (GIP) are important subnets for constructing the piRNA similarity network. These two similarity network are described in detail below. As a carrier of the genetic information and functions of RNA, sequences play a decisive role in many processes such as translation and catalysis. ncRNA silences or degrades other RNAs through complementary base pairing. Here, the *k*-mer algorithm is utilized to quantify the similarity of the sequences [34]. The core idea of the algorithm is to count the number of subsequences in the sequence (such as AAA, ACG, UCG, etc.) and calculate its Z-score. The calculation process is as follows:

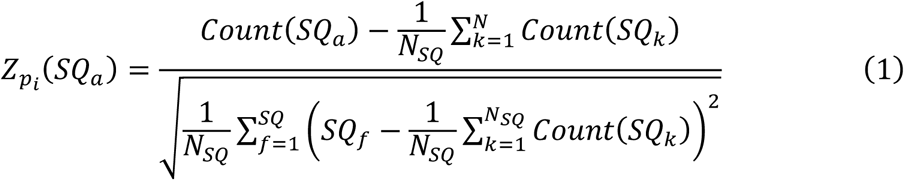

where *p*_*i*_ is the *i*-th piRNA. *SQ*_*a*_ is the *a*-th subsequence. *N*_*SQ*_ is the number of subsequences. *Count*(*SQ*_*a*_) is the number of *SQ*_*a*_. Therefore, the sequence similarity *PS*_*seq*_ between piRNAs can be calculated:

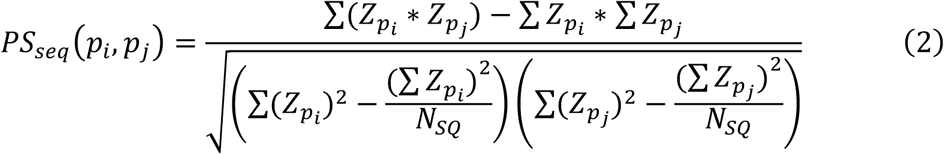

Then, the piRNA gaussian interaction profile kernel similarity can be calculated from the piRNA-disease association network. Specifically, the gaussian interaction profile kernel similarity *PS*_*guss*_ between piRNA *p*_*i*_ and piRNA *p*_*j*_ is defined as follows:

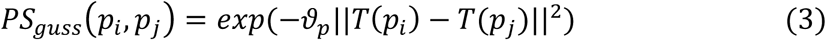

where *T*(*p*_*i*_) is the vector of the relationship between piRNA *p*_*i*_ and all diseases. If piRNA *p*_*i*_ is related to disease *d*, we set the value as 1, otherwise set as 0. *ϑ*_*p*_ is the kernel width coefficient and is defined:

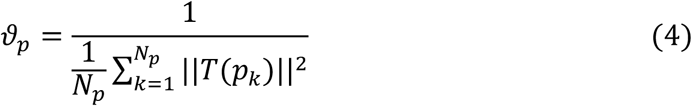

where *N*_*p*_ is the number of piRNAs. According to the piRNA sequence similarity network and the piRNA Gaussian interaction profile kernel similarity network, the similar network of piRNA can be constructed:

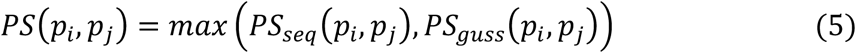

#### The disease similarity network

Disease functional similarity network and disease Gaussian interaction profile kernel similarity network (GIP) are important subnets for constructing disease similarity network. The two similarity networks are described in detail below. At present, there are few methods that can quantify the feature of diseases. Many methods use Medical Subject Headings (MeSH) to construct directed acyclic graphs, and calculate the semantic similarity between two diseases by the relative positions in the directed acyclic graphs [35]. The similarity matrix calculated by this method is very sparse as shown in Figure 1(A) and the coverage is limited. To this end, we propose unified embedding to the disease representation, called Dgene, provides unique 64-dimensional feature vectors for 24166 diseases. In detail, we downloaded 628684 associations about diseases and genes from DisGeNET, http://www.disgenet.org, which contains 24166 diseases and 17543 genes (Table 2). The specific calculation steps are as follows.

**Table 2.** The details of dataset DisGeNET.

| Dataset | Disease | Gene | Association |
| --- | --- | --- | --- |
| DisGeNET | 24166 | 17543 | 628684 |

**Figure 1.**
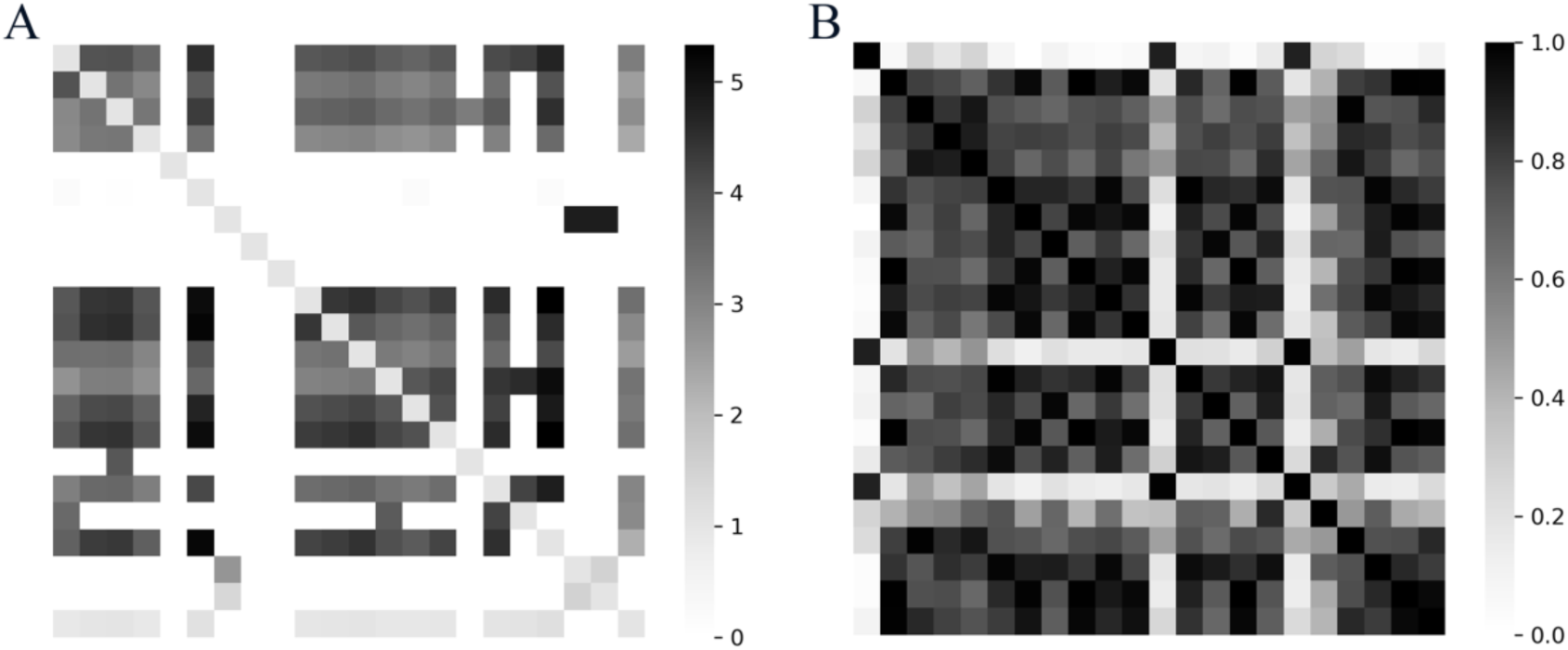
(A) The disease similarity network based on Medical Subject Headings. (B) The disease similarity network based on disease-gene association.

We defined *V*(*d*_*i*_) is the vector of the relationship between disease *d*_*i*_ and all diseases. The *d*_*i*_ is the *i*-th disease. If disease *d*_*i*_ is related to gene *g*, we set the value as 1, otherwise set as 0. Then, the original feature vectors of all diseases are put into a stacked autoencoder to extract deeper abstract information and generate 64-dimensional feature vector *V*’(*d*_*i*_). This operation has the following advantages: reducing the redundancy of the original data and the subsequent calculation costs. According to *V*’(*d*_*i*_), we can calculate the disease functional similarity networ:

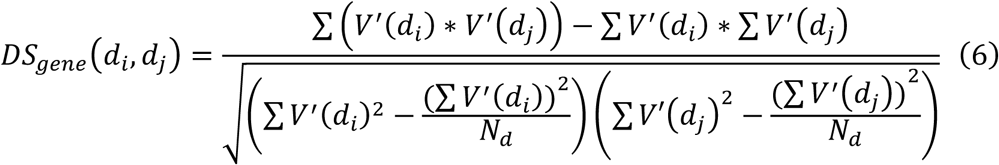

where *N*_*d*_ is the number of diseases in piRD. The visualized disease function similarity matrix network is shown in Figure 1 (B). Compared with disease semantic similarity, Dgene-based disease functional similarity covers more diseases. At the same time, in Dgene, each disease has a unique corresponding 64-dimensional feature vector, and the time-cost of Dgene-based method is much lower than that of MeSH-based method when dealing with different prediction tasks (Table 3).

**Table 3.** Comparison of different types of disease similarity.

| Method | Benchmark dataset | CPU time/pair (seconds)<sup>a</sup> | Size<sup>b</sup> |
| --- | --- | --- | --- |
| MeSH-based | piRD | 378.4498 | 484 |
| Dgene-based | piRD | 0.2919 | 484 |

<sup>a</sup>The average CPU time (seconds) required to calculate the disease similarity matrix.
<sup>b</sup>The size of the disease similarity matrix

Then, the disease gaussian interaction profile kernel similarity can be calculated. In detail, the gaussian interaction profile kernel similarity *DS*_*guss*_ between disease *d*_*i*_ and disease *d*_*j*_ is defined as follows:

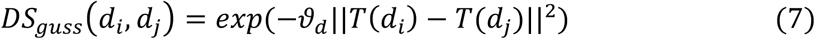

where *T*(*d*_*i*_) is the vector of the relationship between piRNA *d*_*i*_ and all diseases. If disease *d*_*i*_ is related to piRNA *p*, we set the value as 1, otherwise set as 0. *ϑ*_*d*_ is the kernel width coefficient and is defined:

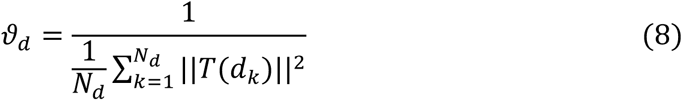

where *N*_*d*_ is the number of disease. According to the disease functional similarity network and the disease Gaussian interaction profile kernel similarity network, the similar network of disease can be figured:

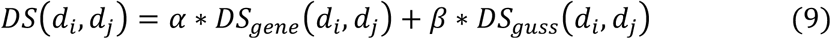

where *α* and *β* are contribution coefficients, and *α* + *β* = 1. Finally, based on the calculated *DS* and *PS*, and the piRNA-disease association network *DP* constructed from the benchmark dataset, the duplex network *Adj* can be defined:

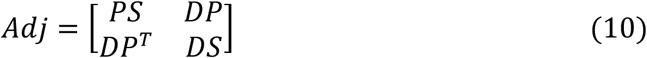

### 2.3 Structural consistency index

In 2015, Liu *et al*. proposed a method for assessing the predictability of complex network links without any prior knowledge of the network, called the “structural consistency” index [36]. Due to its excellent performance and wide versatility in prediction work in other fields[37], this method applied to piRNA-disease duplex networks. In the previous section, we introduced the duplex network *Adj*, which is composed of adjacency matrix *DP*, disease similarity matrix *DS*, and piRNA similarity matrix *PS*. Therefore, the edges in the duplex network composed of piRNA and disease are weighted. The duplex network can be defined as *DN*(*N, E, W*). Where *N* represents the set of nodes composed of piRNA and disease in the network, *E* represents the set of edges in the network, and *W* represents the weight of each edge. The perturbation set *E*^*p*^ consists of randomly selected edges of the duplex network, and the remaining edges *E* − *E*^*p*^ are defined as *E*^*r*^. The duplex network made up of the perturbed set *E*^*p*^ is denoted as *Adj*^*p*^, and the duplex network made up of the remaining edges set *E*^*r*^ is denoted as *Adj*^*r*^. In particular, *Adj* = *Adj*^*p*^ + *Adj*^*r*^. As the duplex network is undirected, *Adj, Adj*^*p*^, and *Adj*^*r*^ are all real symmetric matrices. Therefore, we can diagonalize *Adj*^*r*^ as follow:

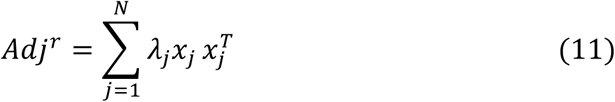

where *λ*_*j*_ is the eigenvalue of the duplex network *Adj*^*r*^, and *x*_*j*_ is the corresponding orthogonal and normalized eigenvector. Then, transforming the perturbation set *E*^*p*^ into a perturbation matrix through first-order approximation. In the case of repeated eigenvalues, there will be degradation eigenvalues. Therefore, we discuss two situations that can arise. The first is the case where there are nonduplicate eigenvalues. After perturbing *Adj*^*r*^, the eigenvalue changes from *λ*_*j*_ to 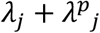, and the corresponding eigenvector changes from *x*_*j*_ to 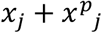 Then, left-multiplying eigenfunction:

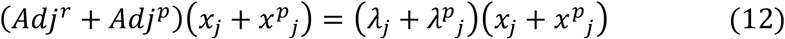

According to the above formula, when the second-order terms 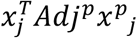 and 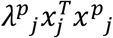 are ignored, we can obtain the following results.

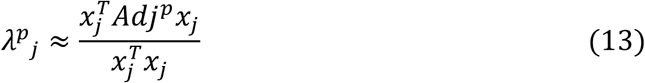

Under the condition that the eigenvector is kept unchanged, the matrix after the perturbation can be represented by the eigenvalue of *Adj*^*p*^corresponding to perturbation set *E*^*p*^ and the eigenvalue of *Adj*^*r*^.

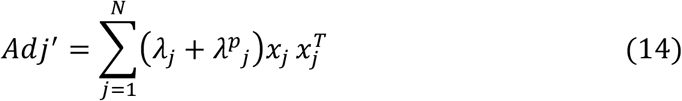

The second is the case where there are duplicate eigenvalues. This means that there are multiple eigenvectors corresponding to the same eigenvalue. Therefore, the eigenvalue is defined as *λ*_*ji*_, where *j* refers to the *j*th eigenvalue and *i* refers to the *i*th eigenvector corresponding to the eigenvalue. According to previous research, we assume that any linear combination of eigenvectors corresponding to the same eigenvalue is still a eigenvector [38]. After perturbing *Adj*^*r*^, the symmetry of the node will be improved. For this reason, the degenerate eigenvalues needs to be selected to continuously convert them into nondegenerate eigenvalues [39]. In detail, the selected eigenvectors is defined as 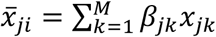. Where *M* represents the number of all eigenvectors corresponding to the same eigenvalue. Then, the eigenfunction can be defined as follows:

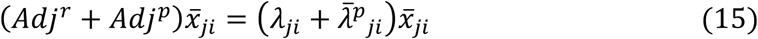

giving us

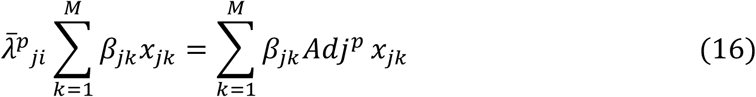

Then, left-multiplying 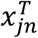 where *n* ∈ [1,. ., n]:

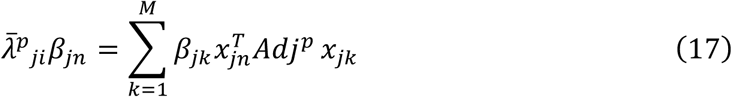

The above equation transformed into matrix form can be written as:

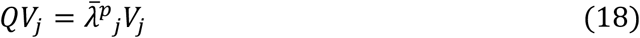

where *Q* is a matrix of size *M* × *M*. The element *Q*_*nk*_ can be calculated by 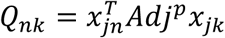. *V*_*j*_ is the column vector*β*_*jk*_. Finally, the matrix after the perturbation *Adj*^’^ can be obtained by replacing and *x*_*j*_ and 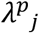 with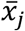 and 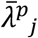 Since the eigenvector can quantify the network structure, if the eigenvectors of *Adj* and*Adj*’ tend to be the same, then the perturbation cannot obviously change the matrix structure, that is, *Adj* has high structural consistency. Structural consistency *τ* can be defined as follows:

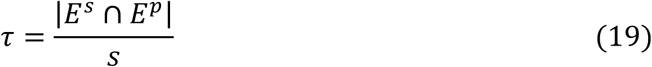

where *s* is the number of edges in *E*^*p*^, and *E*^*s*^ is the *s* edges with the highest predicted score in *Adj*’. In Table 4, we calculated the structural consistency *τ* of the five related networks mentioned above, where *E*^*p*^ is 10% of *E*. It can be seen from the table that the structural consistency scores of DP+PS and duplex network are higher than DP, indicating that including more information can improve the structural consistency of the matrix. There is a gap between DP+MeSH, DP+DS and DP scores, which shows that too little added information will decrease the structural consistency score. In addition, DP+DS has a higher structural consistency score than DP + MeSH, which shows that DP+DS can improve the predictability of the matrix.

**Table 4.** Comparison of the structural consistency of the five networks.

| Network | Structural consistency | Additional information |
| --- | --- | --- |
| $DP$ | $0.3680 \pm 0.0160$ | N/A |
| $DP + PS$ | $0.5900 \pm 0.0331$ | piRNA similarity |
| $DP + MeSH$ | $0.0165 \pm 0.0077$ | Disease similarity |
| $DP + DS$ | $0.3589 \pm 0.0167$ | Disease similarity |
| $Duplex\ network(Adj)$ | $0.6023 \pm 0.0297$ | piRNA and disease similarity |

### 2.4 Method overview

In this paper, a novel method based on duplex network and structural perturbations is proposed to identify biologically significant, yet unmapped piRNA-disease associations, called SPRDA. The SPRDA has three main steps as shown in Figure 2. First, disease function information, piRNA sequence information and original piRNA-disease association information were used to construct the disease functional similarity network *DS*_*gene*_, the piRNA sequence similarity network *PS*_*seq*_, the disease Gaussian interaction profile kernel similarity network*DS*_*guss*_, and the piRNA Gaussian interaction profile kernel similarity network *PS*_*guss*_, respectively. Based on the four similar networks obtained above, the disease similar network *DS* and the piRNA similar network *PS* can be integrated. Second, the duplex network *Adj* is built based on the calculated disease similarity network *DS*, piRNA similarity network *PS*, and known piRNA-disease associations *PD*. Finally, the predicted scores of the unknown associations in the duplex network *Adj* are calculated through structural perturbation.

**Figure 2.**
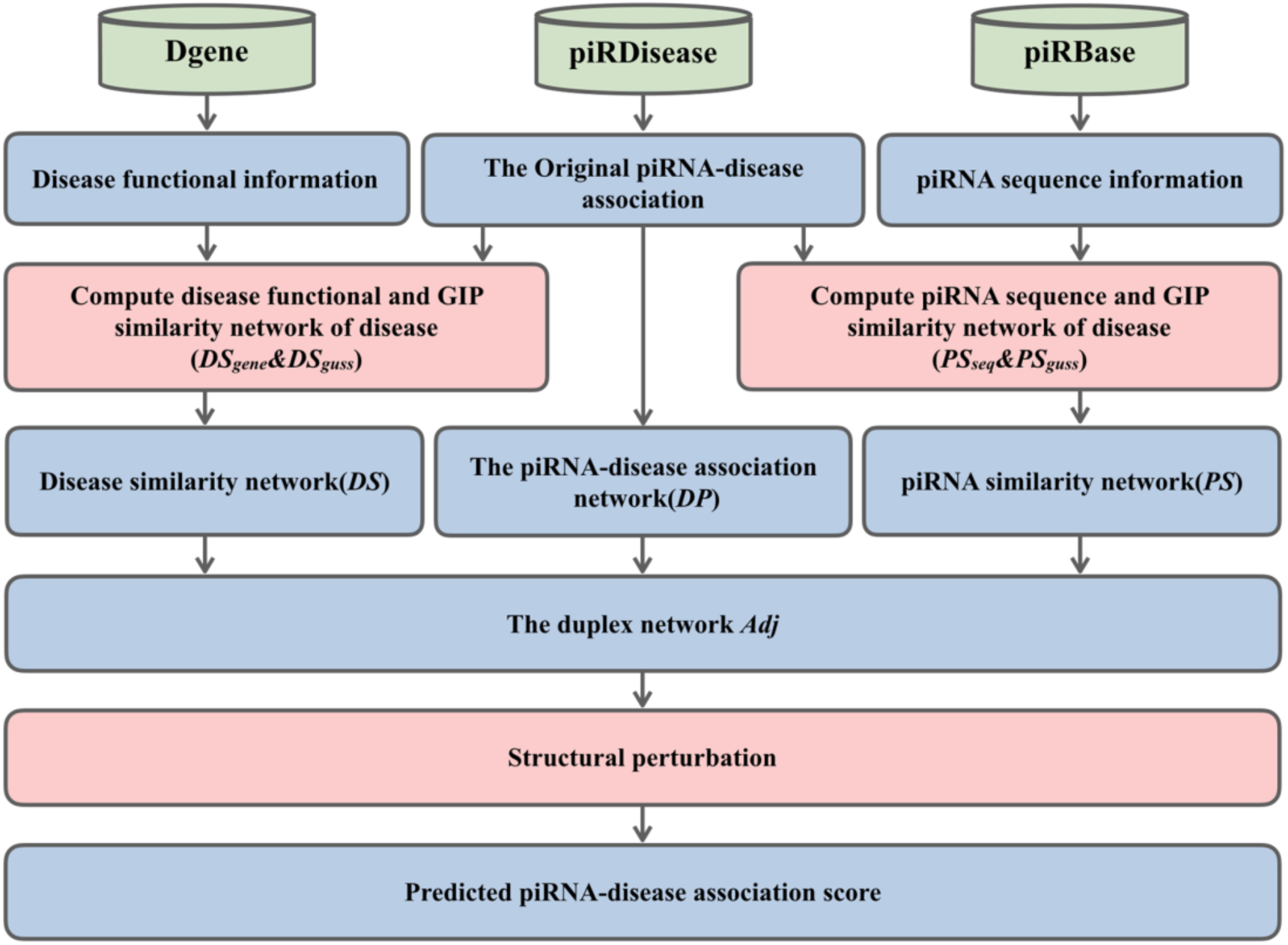
The main step of SPRDA to identify biologically significant, yet unmapped piRNA-disease associations.

## 3 Experimental Results

### 3.1 Parameter optimization

Parameters often affect the performance of the model. To obtain the optimal model, two parameters were used to study its effect on the prediction performance of SPRDA. They are *a* and *t*, where *a* is the percentage of *E*^*p*^ in *E* and *t* is the number of perturbations, that is, the number of times that Equation 14 is executed. In order to more fully evaluate the performance of the model, five-fold cross-validation was used. It is worth noting that this model is different from the traditional machine learning model, so the evaluation index is different. According to previous studies [40], we calculated the evaluation indexes of the model under different precision (*Pre*.). The precision (*Pre*.) is defined as:

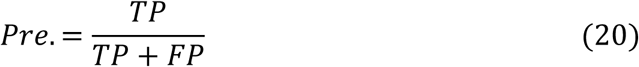

Firstly, we explored the effect of the parameter *t* on the prediction performance of SPRDA based on duplex network when the parameter *a* is fixed at 10%, and shown in Table 5. We can see from the table that as the number of perturbations increases, the various evaluation indexes of the model also increase. In addition, the performance of SPRDA is optimal when *t* = 10, and it stabilizes at *t* = 12. Therefore, in the following experiments, *t* is set to 10. Secondly, the influence of parameter *a* on the prediction performance of SPRDA when the parameter *t* is fixed at 10 is discussed, as shown in Table 6. From the table, we can see that with the increase of parameter a, the various evaluation indexes of the model tend to increase first and then decrease. When *a* = 40%, the performance of SPRDA is optimal. When *a* = 50%, the performance decreases. It is worth noting that when *a* = 30% and a = 40%, the performance of the proposed model is very close, but when *a* = 40%, the calculation time increases significantly. Therefore, in order to balance the efficiency of the model, *a* is set to 30% in the following experiments.

**Table 5.** The effect of the parameter *t* on the performance of SPRDA when fixed parameter *a* = 10%.

| Parameter | *Pre.* | AUC | Accuracy | Recall | MCC |
| --- | --- | --- | --- | --- | --- |
| $t=2$ | 90% | 0.9007±0.0171 | 0.8284±0.0204 | 0.7918±0.0285 | 0.7137±0.0285 |
| | 95% | 0.8969±0.0044 | 0.7987±0.0269 | 0.7331±0.0303 | 0.6657±0.0384 |
| $t=4$ | 90% | 0.9097±0.0127 | 0.8622±0.0108 | 0.8417±0.0173 | 0.7621±0.0161 |
| | 95% | 0.9102±0.0149 | 0.8013±0.0808 | 0.7442±0.0900 | 0.6723±0.1155 |
| $t=6$ | 90% | 0.9117±0.0152 | 0.8556±0.0143 | 0.8316±0.0222 | 0.7525±0.0209 |
| | 95% | 0.9157±0.0082 | 0.8322±0.0411 | 0.7751±0.0515 | 0.7152±0.0608 |
| $t=8$ | 90% | 0.9139±0.0091 | 0.8677±0.0324 | 0.8526±0.0498 | 0.7712±0.0472 |
| | 95% | 0.9145±0.0126 | 0.8380±0.0518 | 0.7842±0.0660 | 0.7245±0.0771 |
| **$t=10$** | **90%** | **0.9158±0.0107** | **0.8684±0.0260** | **0.8531±0.0427** | **0.7720±0.0391** |
|  | **95%** | **0.9140±0.0102** | **0.8231±0.0593** | **0.7664±0.0716** | **0.7029±0.0866** |
| $t=12$ | 90% | 0.9153±0.0085 | 0.8680±0.0239 | 0.8521±0.0378 | 0.7713±0.0353 |
| | 95% | 0.9154±0.0091 | 0.8305±0.0355 | 0.7721±0.0420 | 0.7123±0.0514 |

**Table 6.** The effect of the parameter *a* on the performance of SPRDA when fixed parameter *t* = 10.

| Parameter | *Pre.* | AUC | Accuracy | Recall | MCC |
| --- | --- | --- | --- | --- | --- |
| $\alpha=5\%$ | 90% | 0.8904±0.0200 | 0.8437±0.0345 | 0.8155±0.0483 | 0.7361±0.0482 |
| | 95% | 0.8896±0.0128 | 0.7621±0.0675 | 0.6981±0.0681 | 0.6153±0.0941 |
| $\alpha=10\%$ | 90% | 0.9158±0.0107 | 0.8684±0.0260 | 0.8531±0.0427 | 0.7720±0.0391 |
| | 95% | 0.9140±0.0102 | 0.8231±0.0593 | 0.7664±0.0716 | 0.7029±0.0866 |
| $\alpha=20\%$ | 90% | 0.9293±0.0112 | 0.8792±0.0274 | 0.8712±0.0458 | 0.7884±0.0416 |
| | 95% | 0.9433±0.0092 | 0.8726±0.0464 | 0.8292±0.0464 | 0.7765±0.0509 |
| $\alpha=30\%$ | 90% | **0.9524±0.0085** | **0.9002±0.0183** | **0.9073±0.0335** | **0.8207±0.0290** |
|  | 95% | **0.9529±0.0088** | **0.8734±0.0452** | **0.8325±0.0671** | **0.7788±0.0715** |
| $\alpha=40\%$ | 90% | 0.9566±0.0080 | 0.9064±0.0166 | 0.9187±0.0313 | 0.8306±0.0267 |
| | 95% | 0.9563±0.0074 | 0.8936±0.0329 | 0.8612±0.0507 | 0.8101±0.0528 |
| $\alpha=50\%$ | 90% | 0.9587±0.0089 | 0.9171±0.0120 | 0.9389±0.0237 | 0.8480±0.0198 |
| | 95% | 0.9589±0.0091 | 0.8792±0.0480 | 0.8415±0.0688 | 0.7881±0.0748 |

### 3.2 The performance of SPRDA in predicting piRNA-disease associations on piRDisease

According to the parameter optimization in the previous section, in this part, the parameter a and t of the experimental model are 30% and 10. As shown in Figure 3, when Pre. = 90%, the average predicted AUC of SPRDA is 0.9524 +/- 0.0085. The AUCs for the five experiments were 0.9603, 0.9493, 0.9624, 0.9435, and 0.9460. When *Pre*. = 95%, the average predicted AUC of SPRDA is 0.9529 +/- 0.0088. The AUCs of the five experiments were 0.9609, 0.9549, 0.9610, 0.9456, and 0.9419. Table 7 lists that when *Pre*. = 90%, the average accuracy, recall, Precision, and Matthews correlation coefficient (MCC) are 0.9002, 0.9073, 0.8927, and 0.8207, respectively. Meanwhile, when *Pre*. = 95%, the average accuracy, recall, accuracy, and Matthews correlation coefficient (MCC) were 0.8734, 0.8325, 0.9422, and 0.7788, respectively. From the above evaluation indexes, it is not hard to see that the outstanding performance of SPRDA may promote research in the field of piRNA.

**Table 7.**
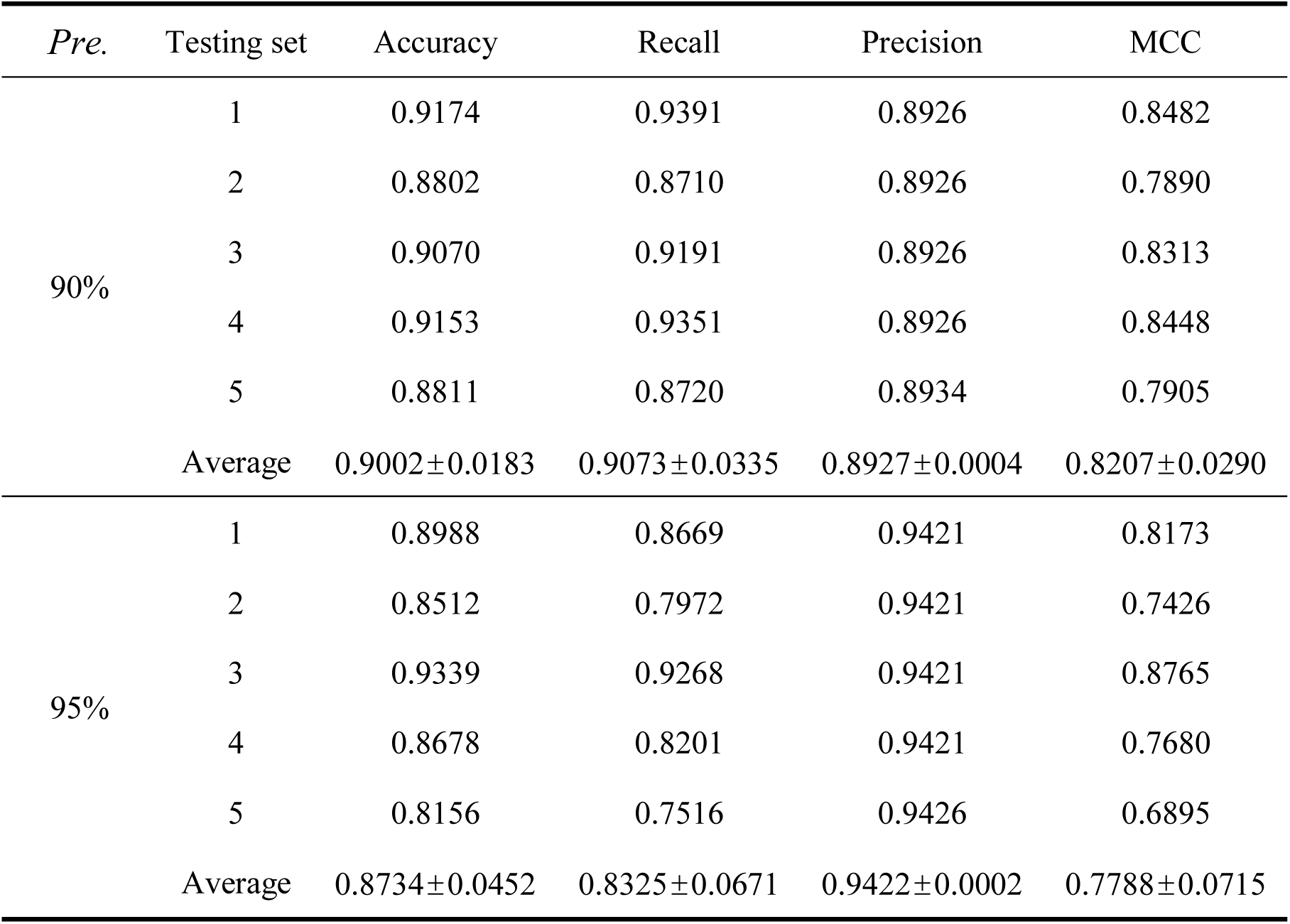
Five-fold cross-validation results performed by SPRDA on piRD dataset.

| *Pre.* | Testing set | Accuracy | Recall | Precision | MCC |
| --- | --- | --- | --- | --- | --- |
| 90% | 1 | 0.9174 | 0.9391 | 0.8926 | 0.8482 |
|  | 2 | 0.8802 | 0.8710 | 0.8926 | 0.7890 |
|  | 3 | 0.9070 | 0.9191 | 0.8926 | 0.8313 |
|  | 4 | 0.9153 | 0.9351 | 0.8926 | 0.8448 |
|  | 5 | 0.8811 | 0.8720 | 0.8934 | 0.7905 |
|  | Average | 0.9002±0.0183 | 0.9073±0.0335 | 0.8927±0.0004 | 0.8207±0.0290 |
| 95% | 1 | 0.8988 | 0.8669 | 0.9421 | 0.8173 |
|  | 2 | 0.8512 | 0.7972 | 0.9421 | 0.7426 |
|  | 3 | 0.9339 | 0.9268 | 0.9421 | 0.8765 |
|  | 4 | 0.8678 | 0.8201 | 0.9421 | 0.7680 |
|  | 5 | 0.8156 | 0.7516 | 0.9426 | 0.6895 |
|  | Average | 0.8734±0.0452 | 0.8325±0.0671 | 0.9422±0.0002 | 0.7788±0.0715 |

**Figure 3.**
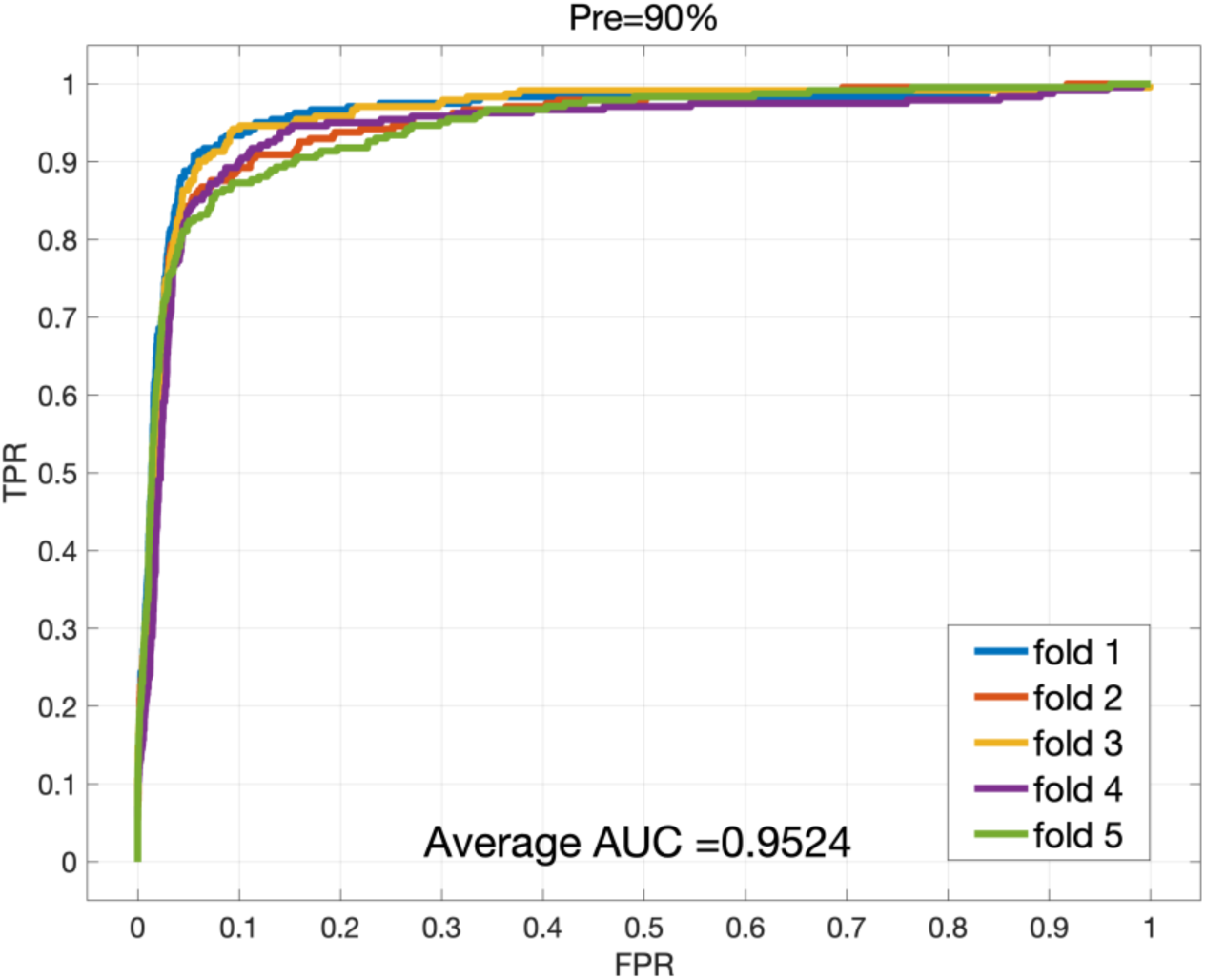
ROC curves performed by SPRDA on piRD (*Pre*.=90%)

### 3.3 Comparison with other four related networks

In Section 2.3, the structural consistency of five related networks is compared. In order to compare these networks more comprehensively, in this section, we compare their performance in predicting piRNA-disease associations. It can be seen from Table 8 that the performance of these related networks tends to be consistent with the structural consistencies. The AUCs of DP+MeSH network and DP+DS network are lower than DP network, which indicates that too little information contained in DS and MeSH reduces the structural consistency, which leads to differences in their AUC. In addition, since the DP+DS network has a higher structural consistency than the DP+MeSH network, the DP+DS network has relatively better prediction performance. what’s more, the duplex network we build has the highest structural consistency due to the integration of information from multiple networks. Similarly, its performance is the highest among the five related networks. This shows that adding DS and PS can effectively improve the prediction performance of the model.

**Table 8.** The AUC across all tested diseases of the four related networks.

| Method | Five-fold cross-validation |  |  |  |  | Average |
| --- | --- | --- | --- | --- | --- | --- |
|  | 1st | 2nd | 3rd | 4th | 5th | (std) |
| *DP* | 0.7955 | 0.8364 | 0.8004 | 0.8256 | 0.7808 | 0.8077<br>(0.0227) |
| *DP+MeSH* | 0.7031 | 0.7014 | 0.6978 | 0.7466 | 0.7479 | 0.7194<br>(0.0255) |
| *DP+DS* | 0.7624 | 0.7768 | 0.7909 | 0.7487 | 0.7659 | 0.7689<br>(0.0158) |
| *DP+PS* | 0.9541 | 0.9413 | 0.9626 | 0.9413 | 0.9479 | 0.9495<br>(0.0091) |
| *Duplex network(Adj)* | 0.9603 | 0.9493 | 0.9624 | 0.9435 | 0.9460 | 0.9523<br>(0.0085) |

**Table 9.** SPRDA predicts performance at 10 disease-specific associations and the number of associations for each disease on the benchmark dataset piRD.

| Disease | Number of associations | AUC |
| --- | --- | --- |
| Alzheimer disease | 221 | 0.8654 |
| Ovarian cancer | 139 | 0.9034 |
| Gastric cancer (recurrence) | 89 | 0.8551 |
| Male infertility | 61 | 0.8382 |
| Breast cancer | 60 | 0.8625 |
| Dysplastic liver nodules and Hepatocellular carcinoma | 58 | 0.7825 |
| Rheumatoid arthritis | 52 | 0.9267 |
| Heart stroke | 42 | 0.827 |
| Head and neck (squamous cell) carcinoma | 37 | 0.9474 |
| Endometrial carcinogenesis | 10 | 0.8548 |

### 3.4 Performance for predicting disease specific associations

In order to verify that SPRDA has sufficient recognition ability in predicting piRNAs corresponding to a single disease, we separately predicted piRNAs related to ten diseases. Specifically, when predicting the specific disease, all associations related to the disease in the benchmark matrix are removed, and the remaining associations are used to construct a duplex network for prediction. In Table 8, we sort these ten diseases from high to low in number. It is not difficult to see that when the proposed model predicts 10 specific disease-related piRNAs, most AUCs are higher than 0.80. In detail, Figure 5 lists the ROC curves and recalls for three diseases. As k increases, recall rises. This phenomenon indicates that the higher the score, the more likely there is a association. The above experiment shows that the proposed model has sufficient recognition ability in this task.

**Figure 4.**
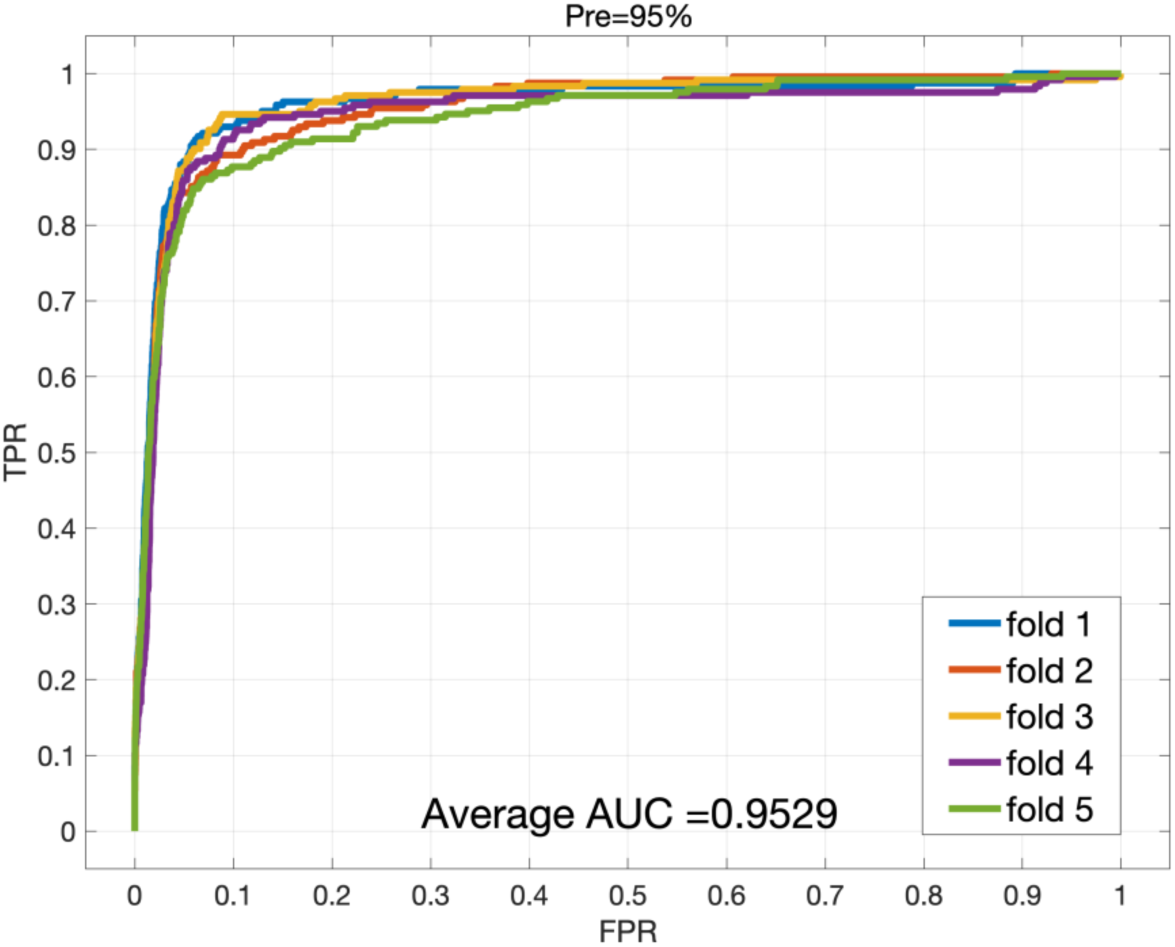
ROC curves performed by SPRDA on piRD (*Pre*.=95%)

**Figure 4.**
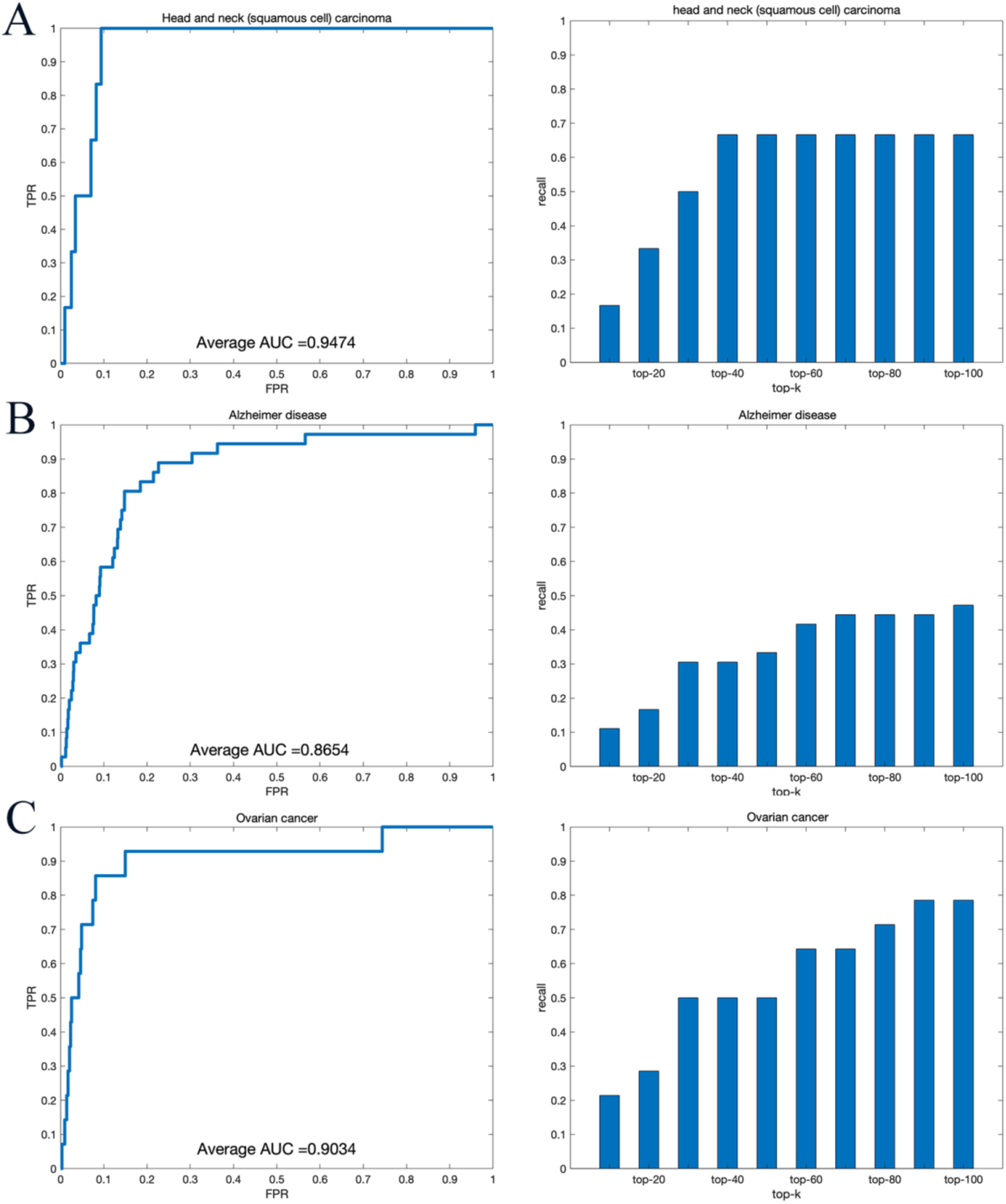
ROC curves and recall for 3 diseases performed by SPRDA on piRD. (A) The ROC curve and recall for Head and neck (squamous cell) carcinoma. (B) The ROC curve and recall for Alzheimer disease. (C) The ROC curve and recall for Ovarian cancer.

## 4 Conclusion

In this study, we proposed a piRNA-disease association prediction based on the structural perturbation method to predict biologically significant, yet unmapped piRNA-disease associations. In particular, we propose a novel disease characterization method (Dgene) and use it to quantify similarity. In detail, the duplex network is constructed based on the piRNA similarity network and disease similarity network extracted from piRNA sequence information, Gaussian interaction profile kernel similarity information and gene-disease association information. Then the structure perturbation method is utilized to predict potential associations on the duplex network. In terms of structural consistency, the duplex network is more predictable than other network structures.

The duplex network and Dgene not only offers excellent prediction performance, but also reduce the time needed for calculation. Experimental results are shown the performance and efficacy of SPRDA in predicting piRNA-disease associations. In the five-fold cross-validation, SPRDA’s AUC, ACC and MCC are 0.9529, 0.8734 and 0.7788, respectively (pre=0.95). To further evaluate the robustness of the proposed model, we tested its performance in predicting 10 disease-specific associations, most AUCs are higher than 0.8000. Overall, the proposed method can provide unique insights into the mechanisms of disease and will further the field of tumor diagnosis and treatment.

## Acknowledgements

This work is supported is supported in part by Awardee of the NSFC Excellent Young Scholars Program, under Grants 61722212, in part by the National Science Foundation of China, under Grants 61873212, 61702444, 61572506, in part by the Pioneer Hundred Talents Program of Chinese Academy of Sciences, in part by the Chinese Postdoctoral Science Foundation, under Grant 2019M653804, in part by the West Light Foundation of The Chinese Academy of Sciences, under Grant 2018-XBQNXZ-B-008. The authors would like to thank all anonymous reviewers for their constructive advices.

